# The intrinsic and regulated proteomes of barley seeds in response to fungal infection

**DOI:** 10.1101/445916

**Authors:** Edward D. Kerr, Toan K. Phung, Christopher H. Caboche, Glen P. Fox, Greg J. Platz, Benjamin L. Schulz

## Abstract

Barley is an important cereal grain used for beer brewing, animal feed, and human food consumption. Fungal disease can impact barley production, as it causes substantial yield loss and lowers seed quality. We used sequential window acquisition of all theoretical ions mass spectrometry (SWATH-MS) to measure and quantify the relative abundance of proteins within seeds of different barley varieties under various fungal pathogen burdens (ProteomeXchange Datasets PXD011303 and PXD014093). Fungal burden in the leaves and stems of barley resulted in changes to the seed proteome. However, these changes were minimal and showed substantial variation among barley samples infected with different pathogens. The limited effect of intrinsic disease resistance on the seed proteome is consistent with the main mediators of disease resistance being present in the leaves and stems of the plant. The seeds of barley varieties accredited for use as malt had higher levels of proteins associated with starch synthesis and beer quality. The proteomic workflows developed and implemented here have potential application in quality control, breeding and processing of barley, and other agricultural products.

## Introduction

Barley is a major cereal grain used as stockfeed for animals, as food for humans, and as the main agricultural product for brewing beer. In 2016, 141 million metric tonnes of barley was produced globally, making it the fourth highest produced cereal commodity behind maize, wheat, and rice [1]. As barley is the main ingredient in brewing, varieties of barley are bred and grown specifically for use as malt in the brewing industry. Many qualities are specifically targeted when breeding and growing malting barley, including high yields, disease resistance, diastase production (starch degrading enzymes), and low levels of β-glucan [2]. In addition, different varieties of barley are grown in Australia for export or domestic markets. This is because the Australian brewing industry tends to use additional sucrose in fermentation, whereas brewers in export markets tend to use additional sources of starch such as rice which require higher levels of diastase enzymes.

Since barley is usually grown in a largely uncontrolled environment, the harvested grain is likely to exhibit variability in grain size, density, starch content, and proteome. As most of the steps in the process of beer brewing rely on proteins, enzymes, and starch from barley, it is expected that variability in the seed will directly affect the beer brewing process and beer quality in complex ways. Diseases are one of the biggest threats to barley production and grain quality, as they may reduce grain size, alter malting quality, and most importantly lower grain yield [3–6]. It is estimated that diseases cause approximately $252 M of losses per annum in barley production in Australia alone [7]. Net form of net blotch (*Pyrenophora teres* f. *teres*), spot form of net blotch (*P. t*. f. *maculata*), and leaf rust (*Puccinia hordei*) are amongst the most common diseases affecting the yield and quality of Australian barley production.

Net blotch, named after the netting pattern that appears on the leaves of infected barley, can also form spot-like lesions. These two distinct symptoms are deemed to be the result of infection by two subspecies (formae) of *P. teres*: *P. teres* f. *teres* and *P. teres* f. *maculata* [8]. These symptoms lead to the corresponding common names of net form of net blotch and spot form of net blotch. Net form of net blotch may cause yield losses in excess of 50% while spot form of net blotch seldom causes losses above 30% [9]. Net blotches are stubble-borne diseases, with *P. teres* f. *teres* also frequently seed-borne [10]. Infection with either form of net blotch can lead to a reduction in seed size and density and can negatively affect the quality of barley for malt and feed [5]. Leaf rust of barley is a disease that produces small, orange-brown pustules on the leaves and leaf sheaths of infected plants. When actively growing, the pustules produce urediniospores which are replaced by black teliospores as they age [11]. The orange-brown coloration of the urediniospores gives the disease the name leaf rust. All three diseases are air-borne with leaf rust the best adapted for wind dispersal [9]. Yield losses associated with leaf rust can be as high as ~ 62% [7, 11–13]. Like net blotch, leaf rust also negatively affects the quality of the grain by reducing grain weight and grain size [6].

Variability in the barley seed proteome due to diseases, barley variety, and other factors is likely to affect seed quality and downstream process efficiencies, and yet is poorly understood. Previous proteomic studies using 2D SDS-PAGE investigated how fungal disease directly affected local plant physiology either in the leaves or seeds during germination [14, 15]. Investigation of the leaf proteome in leaf rust infection identified changes in carbohydrate metabolism, protein degradation, and defence proteins [14], while infection of germinating barley seeds by Fusarium ear blight (*Fusarium graninearum*), caused an increase in proteins involved in carbohydrate metabolism [15]. The use of proteomic analysis of barley seeds has been proposed to link protein abundance to grain quality and germination efficiency [16, 17]. These previous proteomic analyses, along with proteome studies on beer brewing, largely relied on 2D SDS-PAGE technologies [16–18], with only a select number using shotgun proteomics [19, 20]. Several proteomics studies have identified barley proteins throughout the brewing process and in the finished beer [20–23], highlighting that barley proteins are important contributors to the process of beer production, and suggesting that variability in the barley seed proteome will impact beer production process efficiency and quality.

In this study, we used sequential window acquisition of all theoretical ions mass spectrometry (SWATH-MS) to measure and quantify the relative abundance of proteins within barley seeds. This method allowed us to investigate the variability in the proteome of barley seed due to barley variety and burden of fungal disease.

## Methods

### Diseased barley field trial

The Department of Agriculture and Fisheries (DAF) performed the field trial described here in 2015. The work was funded by the Grains Research and Development Corporation (GRDC) as a component of project DAW00245. Three diseases were investigated, net form of net blotch, spot form of net blotch, and leaf rust. For each disease, six varieties of barley selected to balance resistant and susceptible varieties for that disease were tested with diseased (artificially inoculated and not treated with fungicide) and non-diseased (treated with fungicide) treatments in triplicate. Exact trial information is shown in Table 1. Harvested seeds from all three trials were stored at 12 °C, milled to 0.8 mm with a Laboratory Mill 3100 (Perten) cleaned with pressurised air between samples, and stored in Falcon tubes. Two samples were lost in transit and not included in further analysis.

**Table 1.**
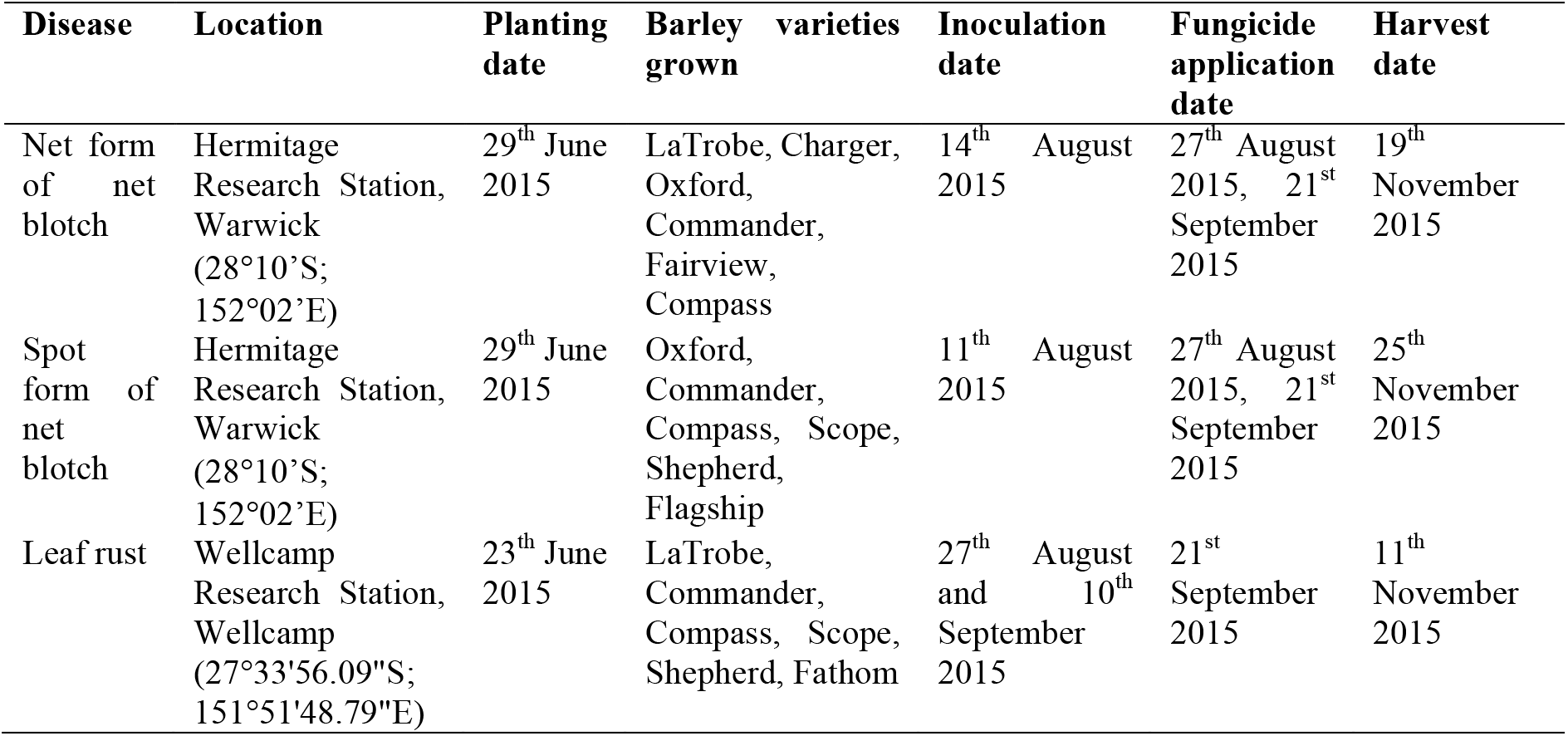
Barley disease field trial information.

### Preparation of milled grain for proteomics

Proteins in milled samples were extracted, denatured, and reduced/alkylated essentially as described [24]. 10 mg milled seed was resuspended in 600 μL of 6 M guanidine hydrochloride, 50 mM Tris HCl buffer pH 8 and 10 mM DTT, and incubated at 30 °C for 30 min with shaking. Cysteines were alkylated by addition of acrylamide to a final concentration of 30 mM and incubation at 30 °C for 1 h with shaking. Excess acrylamide was quenched by addition of DTT to a final concentration of 10 mM and samples were clarified by centrifugation at room temperature at 18,000 rcf for 10 min. To desalt proteins by precipitation, 10 μL of the supernatant was added to 1 mL of 1:1 methanol/acetone and incubated at −20 °C for 16 h. Precipitated proteins were centrifuged at room temperature at 18,000 rcf for 10 min, the supernatant was discarded, and proteins were resuspended in 100 μL of 100 mM ammonium acetate and 1 μg trypsin (Proteomics grade, Sigma). Proteins were digested by incubation at 37 °C for 16 h.

### Mass Spectrometry

Peptides were desalted with C18 ZipTips (Millipore) and measured by LC-ESI-MS/MS using a Prominence nanoLC system (Shimadzu) and TripleTof 5600 instrument with a Nanospray III interface (SCIEX) as previously described [25]. Approximately 1 μg or 0.2 μg desalted peptides, as estimated by ZipTip binding capacity, were injected for data dependent acquisition (DDA) or data independent acquisition (DIA), respectively. LC parameters were identical for DDA and DIA, and LC-MS/MS was performed essentially as previously described [26]. Peptides were separated with buffer A (1% acetonitrile and 0.1% formic acid) and buffer B (80% acetonitrile with 0.1% formic acid) with a gradient of 10–60% buffer B over 14 min, for a total run time of 24 min per sample. Gas and voltage setting were adjusted as required. For DDA analyses, an MS TOF scan from *m*/*z* of 350–1800 was performed for 0.5 s followed by DDA of MS/MS in high sensitivity mode with automated CE selection of the top 20 peptides from *m*/*z* of 40–1800 for 0.05 s per spectrum and dynamic exclusion of peptides for 5 s after 2 selections. Identical LC conditions were used for DIA SWATH, with an MS-TOF scan from an *m*/*z* of 350–1800 for 0.05 s followed by high-sensitivity DIA of MS/MS from *m*/*z* of 50-1800 with 26 *m*/*z* isolation windows with 1 *m*/*z* window overlap each for 0.1 s across an *m*/*z* range of 400–1250. Collision energy was automatically assigned by the Analyst software (SCIEX) based on *m*/*z* window ranges. For validation with SWAT [27] pseudo-PRM, selected milled grain samples (Table S7) were solubilised, digested, and desalted as described above. Peptides were analysed by DIA SWAT with identical LC parameters as above, and with targeted measurement of selected peptide ions (Table S8) with an MS-TOF scan from an *m*/*z* of 350–1800 for 0.05 s followed by high-sensitivity DIA of MS/MS from *m*/*z* of 50-1800 each for 0.1 s.

### Data analysis

Peptides and proteins were identified using ProteinPilot 5.1 (SCIEX), searching against all eukaryotic proteins in UniProtKB (downloaded 29 Jan 2015; 547351 total entries), with settings: sample type, identification; cysteine alkylation, acrylamide; instrument, TripleTof 5600; species, none; ID focus, biological modifications; enzyme, trypsin; search effort, thorough ID. The results from ProteinPilot were used as an ion library to measure the abundance of peptides and proteins using PeakView 2.1 (SCIEX), with settings: shared peptides, allowed; peptide confidence threshold, 99%; false discovery rate, 1%; XIC extraction window, 6 min; XIC width, 75 ppm. The mass spectrometry proteomics data have been deposited to the ProteomeXchange Consortium via the PRIDE [28] partner repository with the dataset identifiers PXD011303 and PXD014093. For protein-centric analyses, protein abundances were normalised to the sum of all protein intensities in a sample. Peakview output was reformatted with a python script (https://github.com/bschulzlab/reformatMS and Supplementary Material – ReformatMS), applying a peptide FDR cut-off of 1% to remove low quality ion measurements for that peptide from each sample, and reformatting appropriate for use with MSstats. Protein abundance differences were determined using MSstats (2.4) in R [29], with Benjamini and Hochberg corrections to adjust for multiple comparisons, and a significance threshold of P = 10^−5^. Gene ontology (GO) term enrichment was determined using GOstats (2.39.1) in R [30] with a significance threshold of P = 0.05. Principal component analysis (PCA) was performed using Python, the machine learning library Scikit-learn (0.19.1), and the data visualisation package Plotly (1.12.2). For comparing malt to feed, varieties were designated as either malt or feed based on various Australian government classification (Table S1) [31–33].

## Results

We aimed to investigate how growth environment, pathogen burden, and barley variety affected malt quality by modifying the molecular composition of barley seeds. To study how these factors affected barley seed proteomes, we performed SWATH-MS proteomics on barley grain grown in a field trial in south east Queensland, Australia, in 2015. Barley was grown in three locations, and at each location the barley plants were infected with a single fungal pathogen: net form of net blotch, spot form of net blotch, or leaf rust, and then either treated or not treated with fungicide (Table 1). As barley at each location was infected with a single disease, location and disease were not separable variables in this study design. Six varieties of barley with varied intrinsic disease resistance were grown at each location (Table S2). Importantly, barley plants were infected in their leaves and stems – the natural sites of infection; and we studied the proteome of the barley seed – the industrially relevant tissue.

### Disease burden results in diverse proteomic responses

Proteins from milled barley seeds were extracted, denatured, reduced/alkylated, precipitated, digested by trypsin, and identified by DDA LC-MS/MS. A total of 220 unique proteins were identified by ProteinPilot (Table S3). We then used SWATH-MS to measure the relative abundance of each protein within each sample quantifying 168 proteins by PeakView with an FDR cutoff of 1%. A rapid LC total method time of 24 min was used to lower instrument time, decrease variability, and increase feasible sample number. We initially used PCA to provide an overview of the proteomic variability in the entire sample set (Fig. 1). This analysis suggested that growth environment / disease was an important factor controlling the proteome of barley seeds, as partial clustering was visible based on location (Fig. 1A and B). No obvious further clustering was apparent within locations when samples were partitioned by pathogen burden (Fig. 1C and 1D).

**Figure 1.**
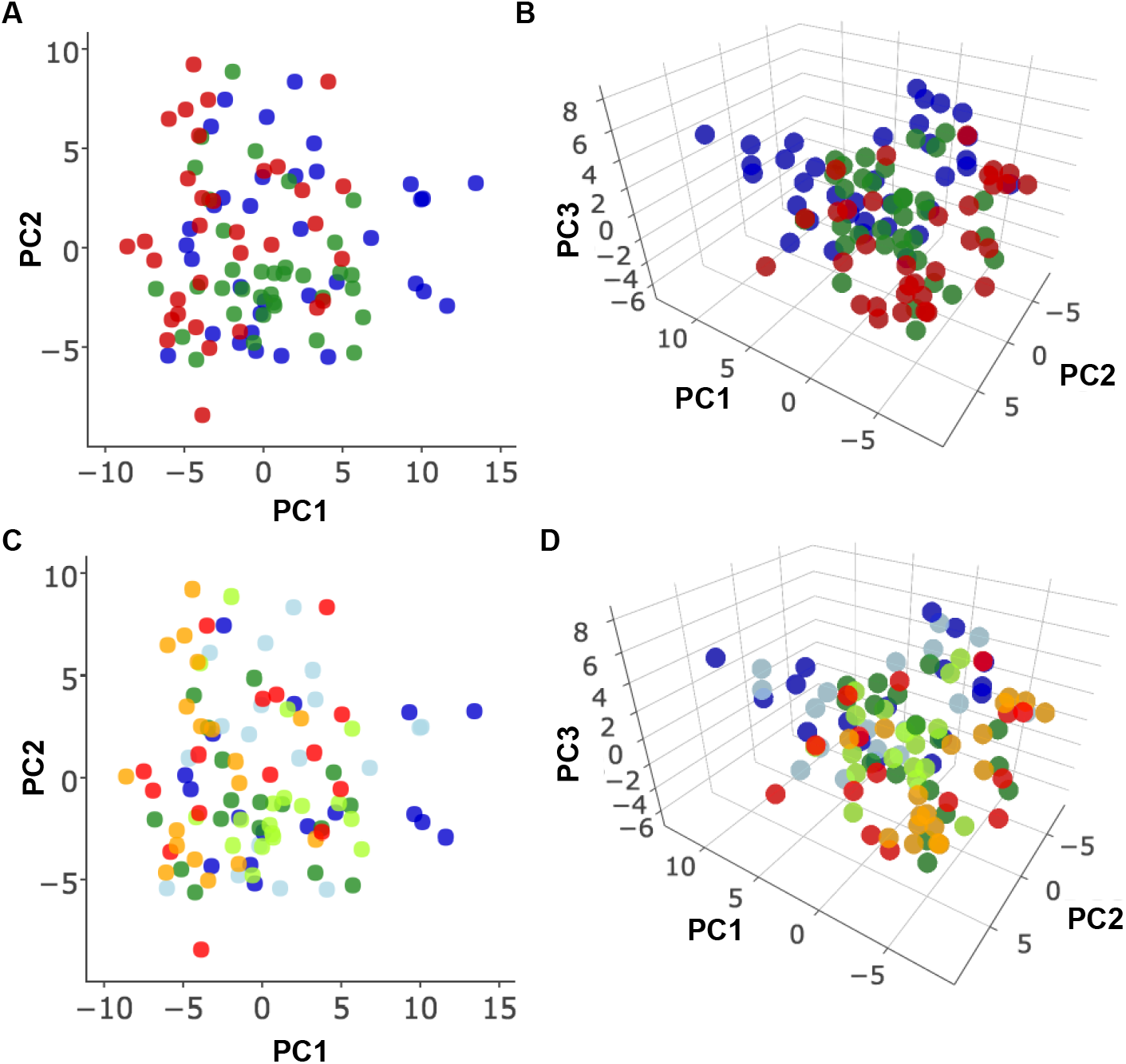
Principle component analysis of the entire dataset highlighting proteomic variation due to disease/location and pathogen burden. **(A)** 2D PCA and **(B)** 3D PCA, both coloured by disease: net form of net blotch (blue), spot form of net blotch (green), and leaf rust (red). **(C)** 2D PCA and **D)** 3D PCA, both coloured by pathogen burden: net form of net blotch – non-diseased (dark blue), net form of net blotch – diseased (light blue), spot form of net blotch – non-diseased (dark green), spot form of net blotch – diseased (light green), leaf rust – non-diseased (red), and leaf rust – diseased (orange). The first component (x-axis) accounted for 12.53% of the total variance, the second 9.02%, and the third an additional 6.78%.

To investigate the effect of pathogen burden on the barley seed proteome we directly compared the proteomes of diseased and non-diseased samples independently for each location/disease, pooling all barley varieties per disease (Table S4). This analysis revealed that disease burden significantly affected the abundance of several proteins across the three diseases/locations (Fig. 2). Interestingly, oxalate oxidase 2 (OXO2) was found to be significantly increased in abundance upon infection with all three pathogens, (Fig. 2B, Table S4). OXO2 is involved in the plant stress response and produces hydrogen peroxide in the apoplast.

**Figure 2.**
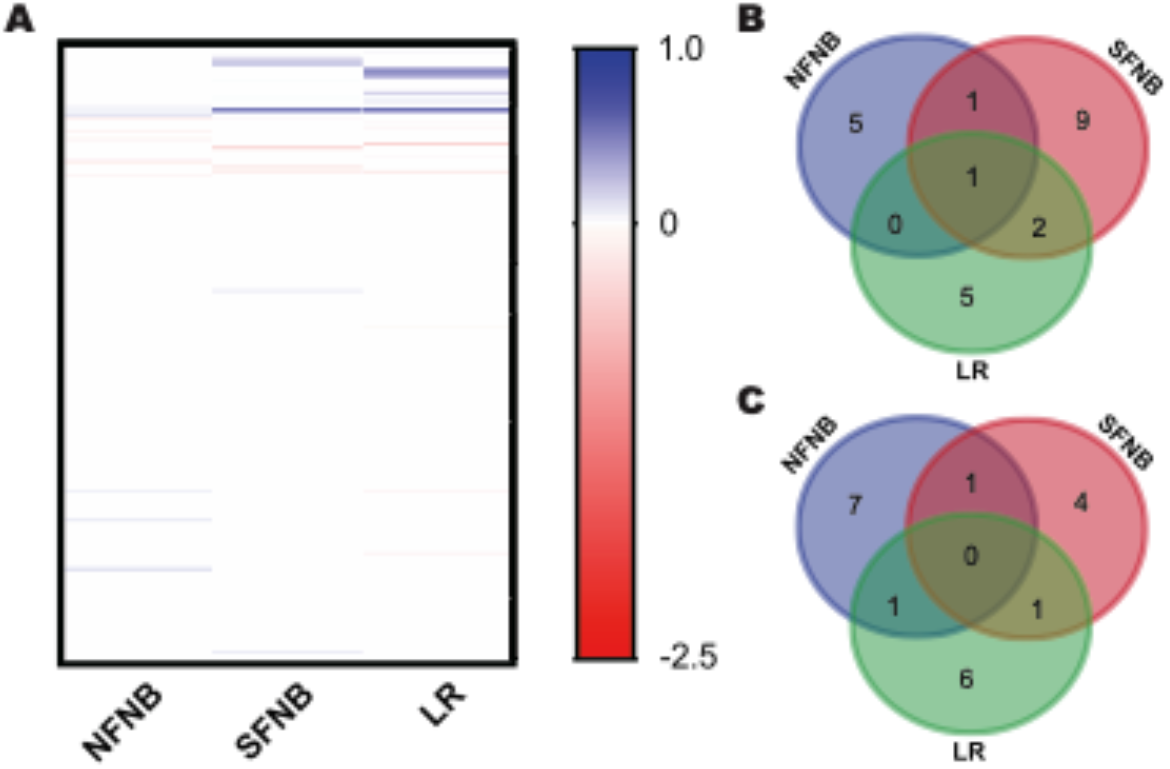
The effect of fungal infection on the barley seed proteome. **(A)** Heat map of significantly differentially abundant proteins between diseased and non-diseased samples. NFNB, net form of net blotch; SFNB, spot form of net blotch; LR, leaf rust. Values shown as log2(fold change) for proteins with significant differences in abundance between diseased and non-diseased (P<10^−5^). Venn Diagram of the number of proteins significantly higher in abundance in **(B)** diseased samples or in **(C)** non-diseased samples.

### Intrinsic disease resistance affects the barley seed proteome

Barley varieties are bred to have specific disease resistance profiles; new varieties are bred and introduced when diseases evolve or become able to infect existing varieties. It is possible that increased disease resistance comes at the cost of productivity or seed quality. We therefore tested if there were differences in the intrinsic proteomes of moderately resistant and susceptible varieties of barley. We independently compared the barley seed proteomes of varieties of barley that were resistant or susceptible to each of the three diseases. Resistance or susceptibility was defined as classified by field trial observations using the GRDC (2016) standard disease resistance rating system (Table S2). We identified a suite of proteins that were significantly different in abundance between moderately resistant and susceptible varieties (Fig. 3 and Table S5). No protein was significantly more abundant in either moderately resistant or susceptible varieties across the three diseases (Fig. 3). However, varieties that were moderately resistant to net form of net blotch showed an overlapping proteomic profile with varieties that were moderately resistant to spot form of net blotch (Fig. 3). In both of these sets of varieties, five proteins were significantly higher in moderately resistant varieties: Sucrose synthase 1 (SUS1), Serpin-Z7 (BSZ7), Gamma-hordein-1 (HOG1), 16.9 kDa class I heat shock protein 1 (HS16A), and Alpha-amylase inhibitor BMAI-1 (IAA1), and seven proteins were significantly higher in susceptible varieties Serpin-Z4 (SPZ4), Alpha-amylase inhibitor BDAI-1 (IAA2), Granule-bound starch synthase 1 (SSG1), Chaperone protein DnaK (DNAK), 1-Cys peroxiredoxin (REHY), endochitinase 1 (CHI1), and Gamma-hordein-3 (HOG3) (Fig. 3B and C).

**Figure 3.**
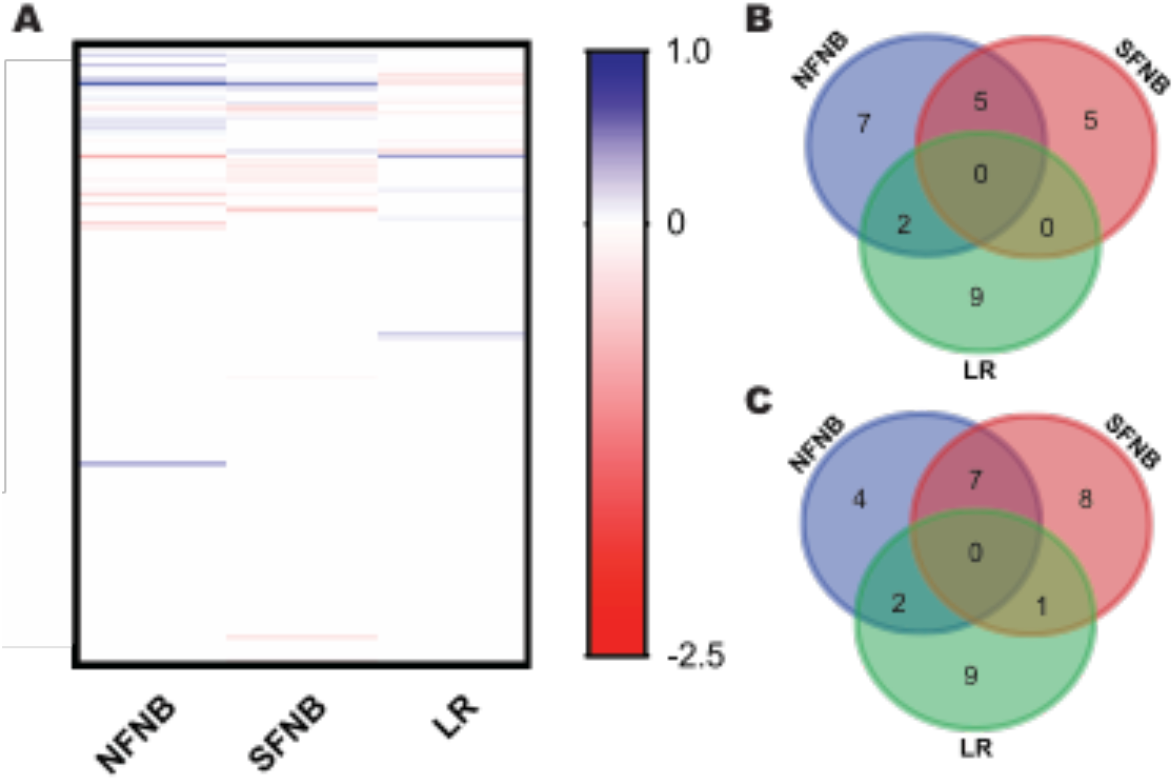
The effect of intrinsic fungal disease resistance on the barley seed proteome. **(A)** Heat map of significantly differentially abundant proteins between moderately resistant and susceptible varieties. NFNB, net form of net blotch; SFNB, spot form of net blotch; LR, leaf rust. Values shown as log_2_(fold change) for proteins with significant differences in abundance between moderately resistant and susceptible varieties (P<10^−5^). Venn Diagram of shared proteins significantly higher in abundance in **(B)** susceptible varieties and in **(C)** moderately resistant varieties.

### Malt quality correlates with specific features of the barley seed proteome

For barley to be used in the brewing process as malt and hence attract a premium price, specific quality measures need to be achieved. To be accredited, a barley variety must be high yielding, adequately disease resistant, and generally perform well in a brewing process. Feed barley, on the other hand, is any variety of barley that has not achieved malt accreditation, and is used for feed for cattle and other livestock. Grain from a malt-accredited variety that is affected by high levels of disease may also be downgraded to feed quality.

We compared the proteomes of barley seeds that had been classified as malt varieties or as feed varieties. Only non-diseased samples were included in this analysis, to remove variability associated with pathogen burden. This analysis showed that 36 proteins were significantly different in abundance between malt and feed varieties (Fig. 4A and Table S6). Of particular interest were several proteins that were more abundant in malt accredited varieties and that have been previously associated with starch synthesis or with beer quality: β-amylase (AMYB), sucrose synthase 1 (SUS1), and sucrose synthase 2 (SUS2) (Fig. 4 and Table S6).

**Figure 4.**
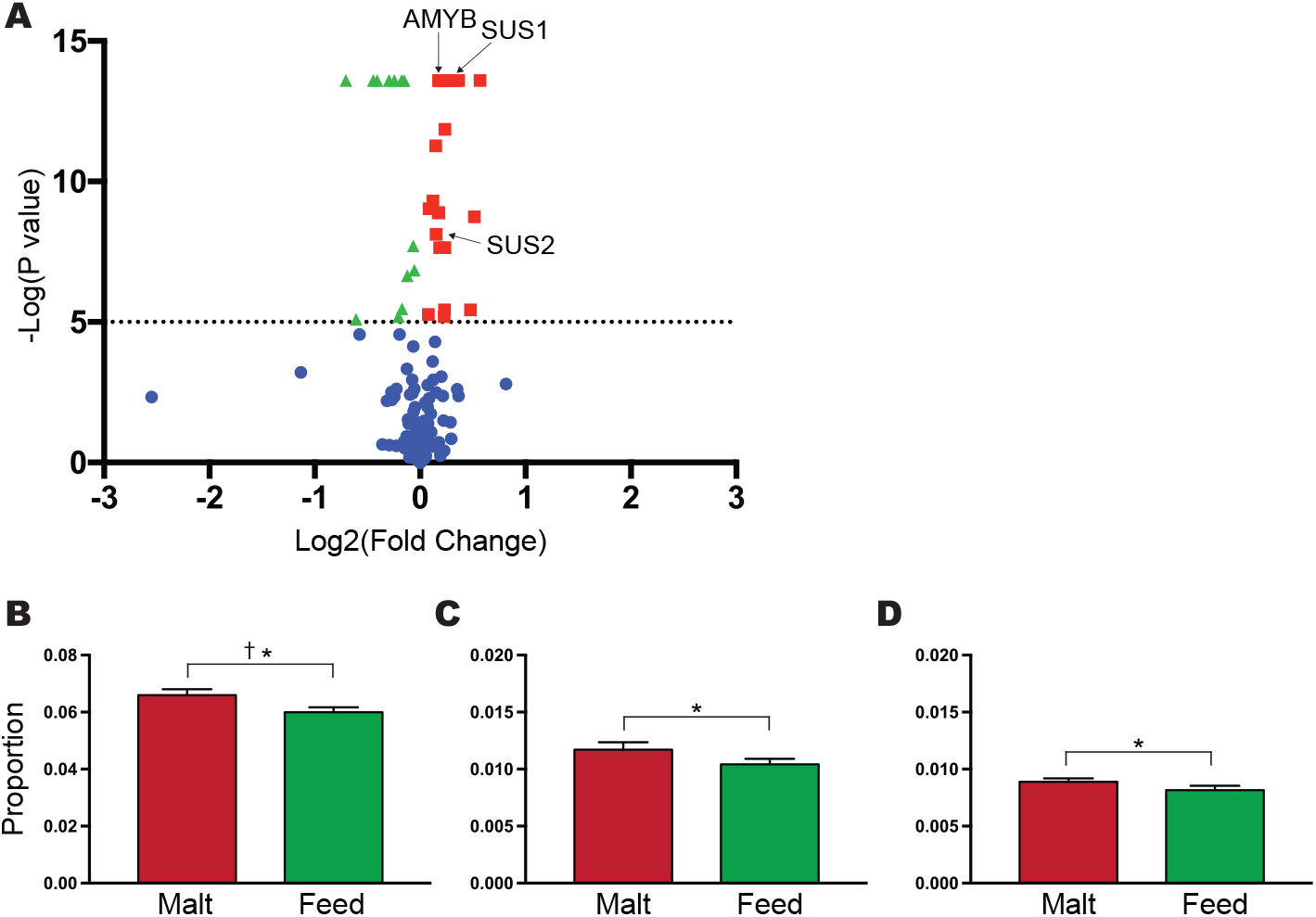
Proteomic features of malt accredited barley. **(A)** Volcano plot of the comparison of the proteome in malt accredited and feed barley. Blue, not significant; red, significantly more abundant in malt varieties; green, significantly more abundant in feed varieties. Abundance of **(B)** β-amylase (AMYB), **(C)** sucrose synthase 1 (SUS1), and **(D)** sucrose synthase 2 (SUS2) in malt (red) and feed (green) varieties. *, P<10^−5^. †, SWAT P<10^−5^.

We validated selected proteomic differences of malt and feed varieties with SWAT, a targeted mass spectrometry approach [27]. We prepared new independent protein extracts from a random subset of samples; digested these with trypsin; used SWAT to measure the abundance of three selected tryptic from each of the three proteins of interest (AMYB, SUS1, and SUS2, as well as housekeeping protein controls phosphoglycerate kinase, cytosolic (PGKY), protein synthesis inhibitor II (RIP2), and enolase 2 (ENO2)); and performed statistical analysis. This validation confirmed that AMYB was significantly more abundant in malt varieties than in feed varieties (Fig. 4B). However, the differences in the abundance of SUS1 and SUS2 detected by SWATH were not validated.

## Discussion

Fungal infection and pathogen burden in the leaves and stem of barley resulted in some changes to the barley seed proteome (Fig. 1 and 2). This suggests that these plants responded systemically to infection, and that fungal infections in the leaves or stem can indeed have an effect on the proteome of the seed. However, in general the changes we observed in the presence of fungal disease were minor and showed considerable variation between barley infected with different pathogens. One protein, OXO2, did show increased abundance in barley infected with all three fungal pathogens (Fig. 2). Oxalate oxidase activity has been previously reported to increase in response to fungal infection of *Erysiphe graminis* f. sp. *hordei* (powdery mildew) in barley leaves [34], but the response in the seed upon pathogen infection has not been previously reported.

The innate resistance status of the barley varieties had only a small effect on the seed proteome (Fig. 3, Table S5). This limited effect of intrinsic disease resistance on the barley seed proteome is consistent with the main mediators of disease resistance being present in the leaves and stems of the plant [35]. That is, disease resistance is likely improved through breeding via altered leaf and stem proteomes, with only subtle changes in the seed. This suggests that breeding for barley disease resistance can be largely independent of seed quality, at least in terms of the proteome. This is valuable information for breeders who can pursue improved disease resistance with confidence that it is unlikely to affect grain quality.

Varieties of barley accredited for use in malting had higher levels of proteins associated with starch synthesis and beer quality [36–38]. β-amylase, non-specific lipid-transfer proteins, and sucrose synthase proteins were all significantly more abundant in malt than feed varieties (Fig. 4 and Table S6). β-amylase (AMYB) is involved in the hydrolysis of starch into fermentable sugars [39–41]. High levels of β-amylase would therefore increase the efficiency of starch degradation during mashing in beer production. Sucrose synthase 1 (SUS1) and sucrose synthase 2 (SUS2) catalyse the conversion of sucrose with nucleotide activated glucose and fructose, and are key regulatory enzymes in the process of starch synthesis [42, 43]. Increased sucrose synthase abundance in malting barley is consistent with selection for high starch content and starch structure more suited to hydrolysis during the malting and brewing process [44]. To our knowledge, this is the first time that proteomic differences have been identified between barley varieties accredited for use in feed or malting. The differences we detected suggest that similar proteomic profiling approaches may be a useful tool in accreditation of malting varieties and in improving the efficiency of variety selection in barley breeding.

PCA revealed that trial location was amongst the largest contributors to the variance in the barley seed proteome. Each geographical condition had many independent variables that could influence barley growth and quality, including soil type and nutrition, level of direct sunlight, temperature, and soil water content [45, 46]. Because of the high level of variability, disentangling the individual contributors to proteomic variation was not possible. In addition, in the experimental design used for the field trial under study, the effects of geographic location could not be separated from fungal disease. Targeted field trial design or use of a more controlled greenhouse setting would allow more detailed investigation of the role of environmental variables in affecting the barley seed proteome.

## Conclusion

Our data provide a glimpse into the molecular complexity and diversity of the barley seed proteome. We provide a high-throughput robust analysis of the response of barley to three important fungal pathogens. Fungal infection and pathogen burden in barley leaves and stems resulted in broad but minimal changes to the barley seed proteome with oxalate oxidase the only protein consistently increased in abundance in infected plants. Accredited malting varieties of barley had significantly higher levels of proteins associated with starch synthesis and beer quality than those classified as feed. More detailed experimentation, possibly under controlled conditions, is required to understand the influence of environment on the barley proteome. The rapid and robust proteomic workflows developed and implemented here have potential application in the quality control of agricultural products, improving breeding efficiency, and studying the manufacturing process of barley and other agricultural products.

## Acknowledgements

We gratefully acknowledge the assistance of Dr Amanda Nouwens and Mr Peter Josh at The University of Queensland School of Chemistry and Molecular Biosciences Mass Spectrometry Facility. Benjamin L. Schulz was funded by an NHMRC Career Development Fellowship APP1087975. Edward D. Kerr was funded by an Advance Queensland PhD Scholarship.

## References

[1] FAOSTAT, United Nations 2016.

[2] Fox, G. P., Panozzo, J. F., Li, C. D., Lance, R. C. M., et al., Molecular basis of barley quality. Australian Journal of Agricultural Research 2003, 54, 1081–1101.

[3] Williams, K., The molecular genetics of disease resistance in barley. Crop and Pasture Science 2003, 54, 1065–1079.

[4] Chełkowski, J., Tyrka, M., Sobkiewicz, A., Resistance genes in barley (Hordeum vulgare L.) and their identification with molecular markers. Journal of Applied Genetics 2003, 44, 291–309.

[5] Grewal, T., Rossnagel, B., Pozniak, C., Scoles, G., Mapping quantitative trait loci associated with barley net blotch resistance. Theoretical and Applied Genetics 2008, 116, 529–539.

[6] Webb, C. A., Fellers, J. P., Cereal rust fungi genomics and the pursuit of virulence and avirulence factors. FEMS Microbiology Letters 2006, 264, 1–7.

[7] Murray, G. M., Brennan, J. P., Estimating disease losses to the Australian barley industry. Australasian Plant Pathology 2010, 39, 85–96.

[8] Smedegård-Petersen, V., Pyrenophora teres f. maculata f. nov. and Pyrenophora teres f. teres on barley in Denmark. Yearbook of the Royal Veterinary and Agricultural University (Copenhagen) 1971, 1971, 124–144.

[9] Department of Agriculture and Fisheries, 2012.

[10] Liu, Z., Ellwood, S. R., Oliver, R. P., Friesen, T. L., Pyrenophora teres: profile of an increasingly damaging barley pathogen. Molecular Plant Pathology 2011, 12, 1–19.

[11] Agriculture Victoria, Department of Economic Development, Jobs, Transport and Resources 2012.

[12] Grain Research and Development Corporation, in: Corporation, G. R. a. D. (Ed.) 2016.

[13] Cotterill, P., Rees, R., Platz, G., Dill-Macky, R., Effects of leaf rust on selected Australian barleys. Australian Journal of Experimental Agriculture 1992, 32, 747–751.

[14] Bernardo, L., Prinsi, B., Negri, A. S., Cattivelli, L., et al., Proteomic characterization of the Rph15 barley resistance gene-mediated defence responses to leaf rust. BMC Genomics 2012, 13, 1.

[15] Yang, F., Svensson, B., Finnie, C., Response of germinating barley seeds to Fusarium graminearum: The first molecular insight into Fusarium seedling blight. Plant Physiology Biochemistry 2011, 49, 1362–1368.

[16] Finnie, C., Melchior, S., Roepstorff, P., Svensson, B., Proteome analysis of grain filling and seed maturation in barley. Plant Physiology 2002, 129, 1308–1319.

[17] Finnie, C., Svensson, B., Barley seed proteomics from spots to structures. Journal of Proteomics 2009, 72, 315–324.

[18] Iimure, T., Nankaku, N., Kihara, M., Yamada, S., Sato, K., Proteome analysis of the wort boiling process. Food Research International 2012, 45, 262–271.

[19] Finnie, C., Svensson, B., Barley seed proteomics from spots to structures. J Proteomics 2009, 72, 315–324.

[20] Mahalingam, R., Shotgun proteomics of the barley seed proteome. BMC Genomics 2017, 18, 44.

[21] Van Nierop, S. N. E., Evans, D. E., Axcell, B. C., Cantrell, I. C., Rautenbach, M., Impact of Different Wort Boiling Temperatures on the Beer Foam Stabilizing Properties of Lipid Transfer Protein 1. Journal of Agricultural and Food Chemistry 2004, 52, 3120–3129.

[22] Schulz, B. L., Phung, T. K., Bruschi, M., Janusz, A., et al., Process Proteomics of Beer Reveals a Dynamic Proteome with Extensive Modifications. Journal of Proteome Research 2018, 17, 1647–1653.

[23] Jégou, S., Douliez, J.-P., Mollé, D., Boivin, P., Marion, D., Evidence of the glycation and denaturation of LTP1 during the malting and brewing process. Journal of Agricultural and Food Chemistry 2001, 49, 4942–4949.

[24] Peak, I. R., Chen, A., Jen, F. E.-C., Jennings, C., et al., Neisseria meningitidis Lacking the Major Porins PorA and PorB Is Viable and Modulates Apoptosis and the Oxidative Burst of Neutrophils. Journal of Proteome Research 2016, 15, 2356–2365.

[25] Xu, Y., Bailey, U. M., Schulz, B. L., Automated measurement of site-specific N-glycosylation occupancy with SWATH-MS. Proteomics 2015, 15, 2177–2186.

[26] Zacchi, L. F., Schulz, B. L., SWATH-MS glycoproteomics reveals consequences of defects in the glycosylation machinery. Molecular & Cellular Proteomics 2016, 15, 2435–2447.

[27] Yeo, K. Y. B., Chrysanthopoulos, P. K., Nouwens, A. S., Marcellin, E., Schulz, B. L., High-performance targeted mass spectrometry with precision data-independent acquisition reveals site-specific glycosylation macroheterogeneity. Analytical Biochemistry 2016, 510, 106–113.

[28] Vizcaíno, J. A., Csordas, A., Del-Toro, N., Dianes, J. A., et al., 2016 update of the PRIDE database and its related tools. Nucleic Acids Research 2016, 44, D447–D456.

[29] Choi, M., Chang, C. Y., Clough, T., Broudy, D., et al., MSstats: an R package for statistical analysis of quantitative mass spectrometry-based proteomic experiments. Bioinformatics 2014, 30, 2524–2526.

[30] Falcon, S., Gentleman, R., Using GOstats to test gene lists for GO term association. Bioinformatics 2007, 23, 257–258.

[31] Agriculture Victoria, Department of Economic Development, Jobs, Transport and Resources 2016.

[32] Department of Agriculture and Fisheries, 2014.

[33] Department of Agriculture and Food, 2016.

[34] Zhang, Z., Collinge, D. B., Thordal-Christensen, H., Germin-like oxalate oxidase, a H2O2-producing enzyme, accumulates in barley attacked by the powdery mildew fungus. The Plant Journal 1995, 8, 139–145.

[35] Breen, S., Williams, S. J., Outram, M., Kobe, B., Solomon, P. S., Emerging Insights into the Functions of Pathogenesis-Related Protein 1. Trends in Plant Science 2017, 22, 871–879.

[36] Sorensen, S., Bech, L., Muldbjerg, M., Beenfeldt, T., Breddam, K., Barley lipid transfer protein 1 is involved in beer foam formation. Technical Quarterly (USA) 1993.

[37] Hao, J., Dong, J., Yu, J., Gu, G., et al., Identification of the major proteins in beer foam by mass spectrometry following sodium dodecyl sulfate-polyacrylamide gel electrophoresis. Journal of the American Society of Brewing Chemists 2006, 64, 166–174.

[38] Evans, D. E., Robinson, L. H., Sheehan, M. C., Tolhurst, R. L., et al., Application of immunological methods to differentiate between foam-positive and haze-active proteins originating from malt. Journal of the American Society of Brewing Chemists 2003, 61, 55–62.

[39] Barth, R., The Chemistry of Beer: The Science in the Suds, John Wiley & Sons 2013.

[40] Briggs, D. E., Boulton, C. A., Brookes, P. A., Stevens, R., Brewing Science and Practice, Woodhead Publishing 2004.

[41] Stewart, G. G., Biochemistry of Foods, Academic Press, San Diego 2013.

[42] Baroja-Fernández, E., Muñoz, F. J., Saikusa, T., Rodríguez-López, M., et al., Sucrose Synthase Catalyzes the de novo Production of ADPglucose Linked to Starch Biosynthesis in Heterotrophic Tissues of Plants. Plant and Cell Physiology 2003, 44, 500–509.

[43] Ghaffari, M. R., Shahinnia, F., Usadel, B., Junker, B., et al., The metabolic signature of biomass formation in barley. Plant and Cell Physiology 2016, 57, 1943–1960.

[44] Chu, S., Hasjim, J., Hickey, L. T., Fox, G., Gilbert, R. G., Structural changes of starch molecules in barley grains during germination. Cereal Chemistry 2014, 91, 431–437.

[45] Finke, P. A., Goense, D., Differences in barley grain yields as a result of soil variability. The Journal of Agricultural Science 1993, 120, 171.

[46] Kemanian, A. R., Stöckle, C. O., Huggins, D. R., Variability of barley radiation-use efficiency. Crop Science 2004, 44, 1662–1672.

