## Supplementary material for "The intrinsic and regulated proteomes of barley seeds in response to fungal infection": readFile.go

```
package fileHandler

import (
	"encoding/csv"
	"io"
	"log"
	"os"
)

type FileObject struct {
	Header     []string
	Filename   string
	OutputChan chan []string
}

func ReadFile(filename string, headerRowNumber int) (fileO FileObject) {
	if headerRowNumber < 1 {
		log.Fatalln("Header Row Number has to be >= 1")
	}
	f, err := os.Open(filename)
	if err != nil {
		log.Fatalln(err)
	}
	reader := csv.NewReader(f)
	reader.Comma = ','
	reader.LazyQuotes = true
	fileO.OutputChan = make(chan []string)
	if headerRowNumber > 1 {
		var headerRows [][]string
		for i := 0; i < headerRowNumber; i++ {
			header, err := reader.Read()
			if err != nil {
				log.Fatalln(err)
			}
			headerRows = append(headerRows, header)
		}
		combinedHeader := make([]string, len(headerRows[0]))
		for i := 0; i < len(combinedHeader); i++ {
			for i2 := 0; i2 < len(headerRows); i2++ {
				combinedHeader[i] += headerRows[i2][i]
			}
		}
		fileO.Header = combinedHeader[:]
	} else {
		fileO.Header, err = reader.Read()
		if err != nil {
			log.Fatalln(err)
		}
	}

	go func() {
		for {
			row, err := reader.Read()
			if err != nil {
				if err == io.EOF {
					break
				}
				log.Fatalln(err)
			}
			fileO.OutputChan <- row
		}
		close(fileO.OutputChan)
		f.Close()
	}()
	return fileO
}
```
