## Supplementary material for "The intrinsic and regulated proteomes of barley seeds in response to fungal infection": userInput.go

```
package input

import (
	"bufio"
	"fmt"
	"os"
	"path/filepath"
	"strings"
	"log"
)

func Input(message string) (filename string, err error) {
	scanner := bufio.NewScanner(os.Stdin)
	fmt.Println(message)
	if scanner.Scan() {
		filename = scanner.Text()
	}
	if err := scanner.Err(); err != nil {
		return "", err
	}
	return filename, nil
}

func Clean(message string) string {
	f, err := filepath.Abs(strings.TrimSuffix(filepath.FromSlash(strings.Replace(message, `"`, "", -1)), " "))
	if err != nil {
		log.Fatalln(err)
	}
	return f
}
```
