## Supplementary material for "The intrinsic and regulated proteomes of barley seeds in response to fungal infection": userInput_test.go

```
package input

import (
	"testing"
	"path/filepath"
)

func TestClean(t *testing.T) {
	result := Clean(`/ ""`)
	f, _ := filepath.Abs(`\`)
	if result != f {
		t.Errorf(`Expected "\" got %v`, result)
	}
}
```
