## Supplementary material for "The intrinsic and regulated proteomes of barley seeds in response to fungal infection": README.md

reformatMS
---

Description:

Uses output from PeakView (an “ions.csv” file and an “FDR.csv” file as a quality filter) and reformats as .csv file appropriate for use in MSstats. 

Appropriate for use with PeakView 2.1 and MSstats 2.4 and above.


Download:

https://github.com/noatgnu/reformatMS/releases

Mac/Linux:

`Downloaded binary needed to be given permission for executing, reading and writing in order for the program to be runnable on a Unix-like system.`

Basic Usage: 
--
Parameter|Function
---|---
-h|Display all available input parameters
-ion|Ion file location in csv format
-fdr|FDR file location in csv format
-out|Output file location in csv format
-t|FDR cutoff threshold (default 0.01)

Example: 

With the script in the same location as inputs file
`.\reformatMS.exe -ion=Ions.csv -fdr=FDR.csv -out=Out.csv -t=0.01`

The user will be prompted to enter each missing parameter besides `-h` and `-t`.

Overall input files rules:
--
- Sample name is by the follow format `{1}_{2}` where `{1}` is the sample title and `{2}` is the sample number.
- Sample columns have to be placed at column number 10 and beyond.
- There must be no blank column within the input files.
- Sample in Ion file input must also be in FDR file.
- Sample column order in Ion file and FDR file must be the same.
