## Supplementary material for "The intrinsic and regulated proteomes of barley seeds in response to fungal infection": reformatMS.go

```
package main

import (
	"flag"
	"github.com/noatgnu/reformatMS/fileHandler"
	"github.com/noatgnu/reformatMS/input"
	"log"
	"os"
	"strconv"
	"strings"
	"bufio"
	"fmt"
)

var swath = flag.String("ion", "", "SWATH Ion File")
var fdr = flag.String("fdr", "", "FDR File")
var out = flag.String("out", "", "Output File")
var threshold = flag.Float64("t", 0.01, "FDR Cutoff threshold")

func init() {
	flag.Parse()
}

func main() {
	openSWATHfile, openFDRfile, filename := TakeUserInput()

	swathFile := fileHandler.ReadFile(openSWATHfile, 1)
	samples := len(swathFile.Header) - 9
	log.Printf("%d Samples", samples)
	fdrFile := fileHandler.ReadFile(openFDRfile, 1)

	fdrMap := ExtractFDRMap(fdrFile, samples)

	o, err := os.Create(filename)
	if err != nil {
		log.Fatalln(err)
	}

	writer := bufio.NewWriter(o)
	writer.WriteString("ProteinName,PeptideSequence,PrecursorCharge,FragmentIon,ProductCharge,IsotopeLabelType,Condition,BioReplicate,Run,Intensity\n")
	outputChan := make(chan string)
	go ProcessIons(outputChan, swathFile, fdrMap, samples)
	for r := range outputChan {
		writer.WriteString(r)
	}
	writer.Flush()
	o.Close()

	log.Println("Completed.")
}

func TakeUserInput() (string, string, string) {
	var openSWATHfile, openFDRfile, filename string
	var err error
	openSWATHfile, err = userInput(openSWATHfile, *swath, "What SWATH-MS file are you opening (written like: SWATH.csv): ", err)
	openSWATHfile = input.Clean(openSWATHfile)
	log.Printf("Input Ion file: %s", openSWATHfile)
	openFDRfile, err = userInput(openFDRfile, *fdr, "What FDR file are you opening (written like: FDR.csv): ", err)
	openFDRfile = input.Clean(openFDRfile)
	log.Printf("Input FDR file: %s", openFDRfile)
	filename, err = userInput(filename, *out, "What are your output file (written like: output.csv): ", err)
	filename = input.Clean(filename)
	log.Printf("Input Output file: %s", filename)
	return openSWATHfile, openFDRfile, filename
}

func ProcessIons(outputChan chan string, swathFile fileHandler.FileObject, fdrMap map[string]map[string][]float64, samples int) {

	//log.Println(fdrMap)
	swathSampleMap := make(map[string][]string)
	log.Println("Processing ions using FDR mapped accession IDs.")
	for c := range swathFile.OutputChan {
		count := 0
		temp := ""
		if v, ok := fdrMap[c[0]]; ok {
			if val, ok := v[c[1]]; ok {
				for i := 0; i < samples; i++ {
					//log.Println(swathFile.Header[9+i])
					var sample []string
					if val1, ok := swathSampleMap[swathFile.Header[9+i]]; ok {
						sample = val1
					} else {
						sample = strings.Split(swathFile.Header[9+i], "_")
						swathSampleMap[swathFile.Header[9+i]] = sample[:]
					}

					row := fmt.Sprintf("%v,%v,%v,%v,%v,%v,%v,%v,%v,", c[0], c[1], c[3], c[7]+c[8], c[6], "L",
						sample[0],
						swathFile.Header[9+i],
						strconv.Itoa(i+1))
					if val[i] < *threshold {
						row += c[9+i]
						if c[9+i] == "" {
							count += 1
						}
					} else {
						row += ""
						count += 1
					}
					row += "\n"
					temp += row

				}
				if count < samples {
					outputChan <- temp

				}
			}
		}

	}
	close(outputChan)
}

func ExtractFDRMap(fdrFile fileHandler.FileObject, samples int) map[string]map[string][]float64 {
	fdrMap := make(map[string]map[string][]float64)
	log.Println("Mapping FDR to accession ID.")
	for c := range fdrFile.OutputChan {
		fdrFail := 0
		if _, ok := fdrMap[c[0]]; !ok {
			fdrMap[c[0]] = make(map[string][]float64)
		}

		var fdrArray []float64
		for i := 0; i < samples; i++ {
			val, err := strconv.ParseFloat(c[7+i], 64)
			if err != nil {
				log.Fatalln(err)
			}

			if val >= *threshold {
				fdrFail++
			}
			fdrArray = append(fdrArray, val)
		}
		if fdrFail < samples {

			fdrMap[c[0]][c[1]] = fdrArray
		}
	}
	return fdrMap
}

func userInput(openSWATHfile string, arg string, message string, err error) (string, error) {
	if arg == "" {
		openSWATHfile, err = input.Input(message)
		if err != nil {
			log.Fatalln(err)
		}
	} else {
		openSWATHfile = arg
	}
	return openSWATHfile, err
}
```
